# Improving neuroimaging headgear placement robustness using facial-landmark guided augmented reality

**DOI:** 10.1101/2025.08.29.660730

**Authors:** Fan-Yu Yen, Yu-An Lin, Qianqian Fang

## Abstract

**Significance:** Accurate and consistent probe placement is crucial in functional near-infrared spectroscopy (fNIRS) and electroencephalogram (EEG) experiments, especially in longitudinal and group-based studies. Both operator experience and subject head shape variability can affect placement accuracy.

**Aim:** We aim to develop an easy-to-use software, NeuroNavigatAR (NNAR), utilizing augmented reality (AR) and machine-learning to estimate and display in real-time the subject’s cranial and head landmarks to guide consistent headgear placement.

**Approach:** By applying a facial recognition toolbox to the image frames extracted from a video camera, we can obtain and continuously track subject-specific three-dimensional (3-D) facial landmarks. Separately, we have precomputed a robust linear transformation between facial landmarks and key cranial landmarks, including nasion and preauricular points, using a large public head-model library consisting of over 1,000 subjects. These allow us to rapidly estmate subject-specific cranial landmarks and subsequently render atlas-derived head landmarks to the subject’s camera stream.

**Results:** An open-source graphical user interface implementing this AR system has achieved a speed of 15 frameper-second using a laptop. A median 10-20 position error of 1.52 cm was found when using a general adult atlas, and is further reduced to 1.33 cm and 0.75 cm when using age-matched atlas models and subject-specific head surfaces, respectively. NNAR demonstrated consistent head-landmark prediction errors across repeated measurement sessions; there is also no statistically significant difference in accuracy across age groups.

**Conclusions:** NNAR is an easy-to-use AR headgear placement monitoring tool that is expected to significantly enhance consistency and reduce setup time for fNIRS and EEG probe donning across a wide range of studies.

## 1 Introduction

Functional near-infrared spectroscopy (fNIRS) and electroencephalogram (EEG) have been widely adopted in neuroimaging studies because they both offer portable instrumentation, high temporal resolution and relatively low system cost. FNIRS employs non-ionizing near-infrared light to measure cortical hemodynamic changes following neuronal activities,^1^ whereas EEG uses electrodes to directly detect neural activation by measuring the electrical potential changes on the scalp.^2^ However, obtaining robust measurement data in both modalities heavily depends on the proper and consistent placement of the probe or headgear,^3^ consisting of optodes in the case of fNIRS and electrodes in the case of EEG. The challenge of consistent placement arises due in part to the variability among subject head shapes and an even greater portion to the operator’s experience.^4^ Inconsistent headgear placement can significantly influence the measurement signal quality, creating challenges when analyzing data from longitudinal and group-based studies.^3^, ^5^

A common approach to guide headgear placement in human experiments typically involves using a head surface-based coordinate system. Systems such as the 10-20, 10-10, or 10-5 systems^6^, ^7^ are defined based on the overall dimensions of the head, determined by several key cranial landmarks. A widely used approach involves manually identifying five cranial landmarks, including nasion (Nz), inion (Iz), left and right preauricular points (LPA, RPA), and vertex (Cz), followed by manually measuring and subdividing the saggital and coronal contour lines into divisions 10% - 20% according to 10-20 system conventions.^7^ Although this method is self-adaptive to the subject’s own head shape and offers good accuracy, performing the measurement carefully could be time-consuming. When measuring these landmarks using a measuring tape, the presence of hairs and the operator’s experience could also result in inconsistencies. An alternate method involves using a pre-fabricated cap with an embedded placement map,^8–12^ usually in the form of optode/electrode mounting grommets or markers. While this approach offers built-in head landmarks to guide the mounting of the optodes/electrodes over the probe, placement errors when donning the headcap over the subject’s head could result in global offsets of all optode/electrode locations, forfeiting some of the accuracy benefits. Another limitation of this approach is that pre-fabricated caps usually come with limited sizes; the 10-20 landmarks on these off-the-shelf headcaps are often based on atlas models and are unable to take into account subject-specific head shapes.^13–16^

To partially compensate for the likely inconsistencies in head cap and probe placement, in many studies, a direct optode/electrode location measurement is often performed as part of the protocol to identify the actual optode/electrode position in a *post hoc* fashion. For example, in some studies, a digitizer is used to measure the three-dimensional (3-D) locations of the cranial and optode/electrode.^17^, ^18^ The digitization step requires additional equipment, taking additional time with added variability due to the uncertainties of human operators. In some other studies, researchers incorporated structural scans of a subject, such as magnetic resonance imaging (MRI), with the head cap or probe labeled with fiducial markers to verify the actual placement.^4^ However, acquiring additional structural images is time consuming and adds additional complexity and cost to the experiment. Regardless, a *post hoc* position measurement could be used to improve data analysis, but fails to provide real-time feedback that can help improving headcap placement prior to measurement.

Over the past decade, the fast development of efficient image processing and computer-vision algorithms offered new tools for researchers to combat the probe placement challenge. Several studies have utilized static or video cameras to guide probe placement during experiment setup. For example, Kawaguchi *et al*. developed two augmented reality (AR) systems using head images and facial landmarks to guide fNIRS probe placement in real-time.^19^, ^20^ While this work offers real-time probe placement guidance, there are a number of limitations: 1) it requires the acquisition of a subject’s own head surface prior to the experiment, 2) image pre-processing steps such as surface extraction and manual labeling of facial landmarks on subject’s head surface are required. In a more recent work^20^ by the same group, the authors addressed some of these limitations by introducing a statistical scalp-cortex correlation atlas, an average brain model from multiple subjects. However, the authors did not provide accuracy assessment, nor provide publicly accessible AR software to be further tested by the community. Song *et al*. proposed a method using AR to guide the EEG electrode placement to reduce the placement error on the same subject across multiple session.^21^ However, this method requires a full head scan during the experiment’s initial setup to capture the subject’s head geometry beforehand. This requires additional cameras and only ensures the following placement is aligned with the initial one, but does not guarantee the placement is aligned with 10-20 system on the head. In addition, Kaewrat *et al*. developed an AR software that overlays EEG electrode positions and a brain model for educational purposes.^22^ While it is effective for visualizing expected electrode locations based on brain anatomy, it is not ideal for experimental setups, as it requires placing fiducial markers on the head beforehand to define the head shape for rendering. Moreover, the system has only been validated on a phantom head and has yet to be tested on human subjects.

In recent years, the rapid emergence of effective image processing approaches based on deeplearning (DL) has demonstrated transformative impact on many research fields, including neuroscience research such as motion tracking or pose analysis for movement disorders ^23^ and emotion recognition.^24^ The availability of many large-sized and well-annotated human facial images and head shape libraries combined with powerful neural network architectures result in dramatically improved accuracy and speed in facial and head shape recognition and recovery, offering exciting new opportunities to build fast and accurate software tools to address the perennial needs of realtime headgear placement guidance and assessment. Open-source facial recognition libraries, such as MediaPipe^25^ developed by Google, have received widespread use among many applications. They are not only capable of extracting and continuously tracking various facial features rapidly and accurately, but also offer excellent robustness, producing accurate results under different viewing angles, lighting conditions, background environments, subject skin-tones, genders, and facial expressions.

In this work, we report an open-source AR tool – NeuroNavigatAR (NNAR) – specifically designed to improve fNIRS/EEG headgear placement accuracy by providing real-time 10-20 landmarks rendered over a video camera stream.^26^ On the one hand, NNAR utilizes the DL-based MediaPipe library to rapidly estimate and continuously track various 3-D facial landmarks based on a subject’s camera image frames. On the other hand, based on a large 3-D head model library, Liverpool–York Head Model (LYHM) dataset,^27^ we have obtained a robust face-to-head landmark transformation with the ability to predict cranial landmark and 10-20 head landmark positions using facial landmarks. Using the LYHM library, we have also systematically characterized the accuracy of our predicted 10-20 head landmark positions by generating synthetic camera images from the textured 3-D head model library at various viewing angles.

In the following Methods section, we first describe the detailed steps of mapping facial landmarks to 10-20 based head landmarks. These steps include creating a library of facial images rendered from a 3-D digital head library at various viewing angles, extracting facial landmarks, manually selecting LPA/RPA to build the ground-truth, and estimating an optimal linear transform mapping from facial landmarks to the expert-determined LPA/RPA positions. With this precomputed linear mapping, we can then render pre-calculated 10-20 positions based on an atlas model to a subject’s camera video frame, creating real-time image overlays. In the Results section, we compare the accuracy of the predicted 10-20 positions between the linear mapping derived from all head models and those derived from subject-specific head models, and between the general adult atlas and the age-matched atlas. Finally, we discuss the limitations of the current work and future steps.

## 2 Method

The overall workflow of the developed headgear placement AR system, NNAR, is illustrated in Fig. 1. The workflow includes two distinct steps. An offline preprocessing step (top) is used to create optimized face-to-head mapping using the LYHM library as well as precompute 10-20 positions from a diverse set of head atlases. The second step (bottom) involves the real-time processing of the facial images captured from a video camera, and apply the pre-computed face-to-head transformation to predict 3-D cranial landmarks such as LPA, RPA, Iz, etc. The real-time predicted subject cranial landmark positions allow us to transform pre-computed 10-20/10-10/10-5 landmarks using a user-specified atlas to the 2-D camera image of the subject and subsequently display/update the 10-20 landmarks as image overlays to guide the operator to position optodes or headgears over the subject’s head.

**Fig 1.**
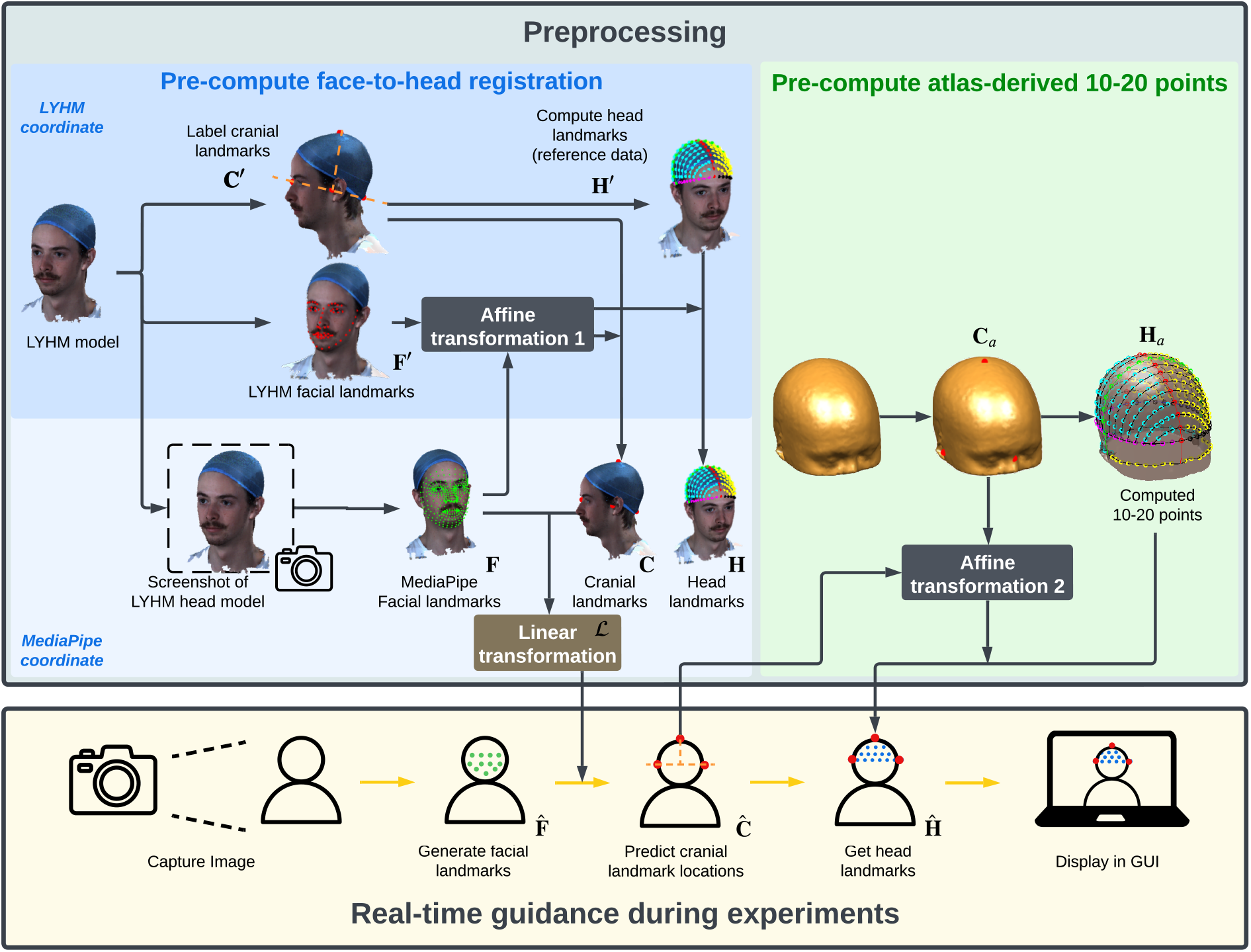
Algorithm workflow of NNAR. The diagram in the upper-left (shaded blue) summarizes the steps to pre-compute the face-to-head landmark mapping using the LYHM head model library. The upper part of this diagram (shaded sky blue) shows the process that is done in the LYHM coordinate system, while the bottom part of this diagram (shaded pale blue) describes the process within the MediaPipe coordinate system. The diagram in the upper-right part (shaded green) describes the steps to pre-compute 10–5 head-landmark positions over a predetermined atlas. The diagram at the bottom (shaded beige) illustrates the processes of real-time computation of the subject-specific head landmarks using a continuous camera image feed and dynamic rendering of the results as overlays on a computer screen. The facial image shown is derived from the LYHM dataset. The dataset authors confirmed that permission to publish these images has been obtained for all individuals in the release.

To ease the description of the involved data processing steps, we use matrix notations **C, F** and **H** to denote the vertically stacked (*x, y, z*) triplets of cranial reference positions (Nz, Iz, LPA, RPA, Cz), facial landmarks and head 10-20 landmarks positions, respectively. The quantities estimated in the atlas are denoted with a subscript of “a”, and those dynamically estimated from a subject’s video frame are denoted with a hat. In addition, 3-D positions stored in the LYHM head model coordinate system are denoted using an apostrophe; those without apostrophe are in the MediaPipe normalized coordinate space. For clarity, the symbolic notation for the respective quantity for each of the processing steps is labeled in Fig. 1.

### 2.1 LYHM head model library and subject inclusion/exclusion criteria

The LYHM dataset contains high-resolution 3-D digitized head models in the form of textured surface models with predefined 68 facial landmarks, covering a total of 1,519 subjects (750 male, 768 female, and 1 transgender), with the subject’s age ranging between 2 and 90 (36.5 *±* 18.2) years old. To test our method over a wide range of head shapes and age groups, we intend to include all head models provided in this library, however, a total of 647 subjects are excluded from this analysis due to one of the following reasons: 1) missing facial texture data (271 subjects), 2) absence of face feature landmarks predefined in the dataset (309 subject; note that 63 subjects are in both categories 1 and 2), 3) unable to locate cranial landmarks due to factors such as hair obstruction, invalid head mesh (e.g., missing parts of the head geometry where cranial landmarks should be located), heads not facing forward which result in the computed Iz being located on the neck or clothing, specifically raised head positions (124 subjects), and 4) unable to compute facial landmarks using MediaPipe (see detail below) (6 subjects). As a result, a total of 872 head models are included in this study, with 465 male and 407 female subjects, and an age of 34.9*±*16.6 years. The included head models contains 808 Caucasians, 27 Asians, 10 Africans, and 37 subjects of other ethnic origins (including 20 mixed-race, 1 Latin/Hispanic and 6 non-disclosed). The LYHM dataset author has confirmed that permission to publish images of individuals in the dataset has been obtained.

For each included 3-D head model, the textured surface mesh is rendered using MATLAB (MathWorks, Natick, MA). We export the 3-D renderings of the head models at 9 different viewing angles as shown in Fig. 2. Each rendering is stored in the PNG (Portable Network Graphic) format with the head region roughly occupying at least around 250×250 pixels.

**Fig 2.**
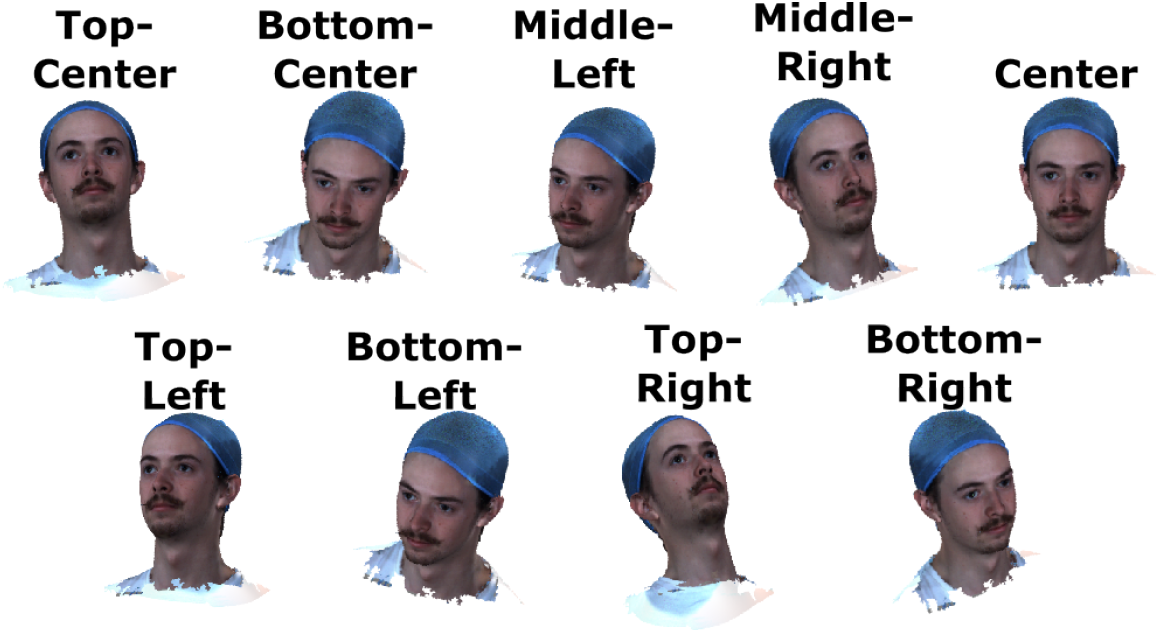
Illustration of the nine distinct head/face orientations used in the face-to-head coordinate system transformation. The screenshots of the 3-D head model are captured from nine viewing angles: three vertical positions (top, middle, and bottom) combined with three horizontal positions (left, center, and right). The facial images are derived from the LYHM dataset, author has confirmed that consent for publication of this specific identity has been obtained.

### 2.2 Facial landmark extraction

We use an open-source library, MediaPipe^25^ – a DL-based facial/gesture recognition framework, to rapidly extract a total of 468 3-D facial landmarks^28^ from each captured 2-D facial image. The extracted facial landmark positions, denoted as **F**, are tessellated in the form of a 3-D triangular surface, covering general facial features such as 1) the overall outline of the face, 2) the outline of the two eyes, 3) the inner and outer contours of the lips, 4) the lower and upper contours of the eyebrows, and 5) the lower edge of the nose contour. We want to particularly highlight that the MediaPipe facial landmark system was primarily developed for automated recognition and processing of facial features. It was not designed for neuroimaging and thus lacked the cranial or head-surface landmarks despite all MediaPipe positions are defined in the 3-D space. From our investigation, the only direct overlap between the MediaPipe-derived facial landmarks and the cranial/head landmark positions is MediaPipe landmark #168, which matches the Nz cranial landmark, i.e. 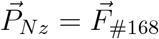.

### 2.3 Manual determination of LPA and RPA cranial positions

To provide reference positions in order to develop an accurate transformation between facial and head landmarks, we have manually selected the left preauricular points 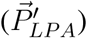 and right preauric-ular points 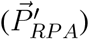 on the 3-D surface of the head model. A subset of the head models was further excluded where the head model had discontinuous or fragmented surfaces around the ears or ear outlines were obscured by hair or headware. We also excluded subjects who lifted their head with an upward gaze, which led to the mislocation of the inion on the subjects’ neck or clothing.

### 2.4 Estimation of Iz and Cz on LYHM head models

Next, we estimate Iz and Cz based on the manually selected LPA/RPA locations on each LYHM head model. This allows us to create the reference cranial landmark positions **C**′ next.

According to the Cleveland Family Study dataset available at the National Sleep Research Resource,^29^ we assume that Nz, LPA, RPA, and Iz are located on the same plane. This allows us to estimate Iz by computing the intersection of the 3-D head surface mesh with a vector extending from 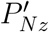 to the midpoint between 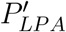 and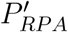. To identify the Cz location 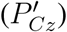 on the head model, we calculate the inner product of the vector from Nz to Iz and the vector from LPA to RPA. The intersection point with the head surface at the top of the head is marked as Cz.

During the process of finding the cranial landmarks, we exclude some subjects due to the incomplete head surface, as previously described in Section 2.1. For each head model, if an open surface is found near the target cranial landmark location, we apply a hole-filling step using Mesh-Fix^30^ and Iso2mesh^31^ to repair the surface. If the mesh fix process is not satisfactory (e.g., hair blocking the cranial landmark locations or the hole is too large to be fixed accurately), we exclude these subjects.

### 2.5 Conversion of cranial reference points from LYHM surface coordinates to MediaPipe normalized coordinate system

The above computed cranial reference points (**C**′) are defined in the LYHM surface coordinate system (in mm units). Next, we transform **C**′ into the MediaPipe normalized coordinate system, in which the coordinates are between 0 and 1. For each set of estimated 3-D MediaPipe facial landmarks **F** from any of the 9 views, we select 10 pairs of matching facial positions that are similarly defined in both the LYHM library (**F**′_(10)_) and the MediaPipe output (**F**_(10)_). The selected positions include the nasion, the corners of the eyes, the tip of the nose, and additional key facial points such as the sides of the nose, the corners of the mouth, and the tip of the chin.

An affine transform is assumed between the selected facial landmarks between the LYHM and MediaPipe coordinate systems in each camera view. The transformation is defined as

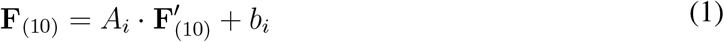

where *A*_*i*_ is a square matrix; *b*_*i*_ is a vector, and *i* = 1, 2, · · ·, 9 is the index of the 9 views.

For each view of each LYHM head model, we compute the *A* and *b* using Eq. 1. This allows us to map the 5 cranial landmarks 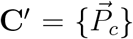 where *c* = *{*Iz, Nz, LPA, RPA, Cz·) computed in Sections 2.3-2.4 on the head model space to the normalized MediaPipe space of the *i*-th view

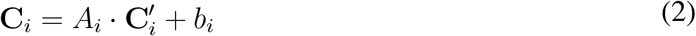

### 2.6 Determination of face-to-head mapping

Our next step is to create a prediction model *ℱ*_*c*_ that transforms the facial landmarks **F**^*i*^ extracted from the *i*-th view of each subject in Section 2.2 to a specific cranial reference point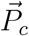, where *c* ∈ *{*Iz, LPA, RPA, Cz·. As a result, for the *i*-th view of every LYHM head model, we have

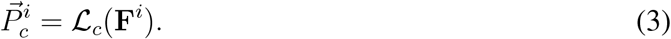

To avoid expensive computation while still capturing major facial features, we select *N*_*f*_ = 20 facial positions from the 468 points provided by MediaPipe. The 20 selected facial landmarks cover key facial features such as face contours, eyes, mouth edges, and nose outlines. Hereinafter, **F** refers to the reduced facial landmark of a size of *N*_*v*_ × (3*N*_*f*_).

With the assumption that each head model is a rigid-body and the prediction model *ℱ*_*c*_ is independent of the camera view, we can concatenate Eq. 3 across all views and form

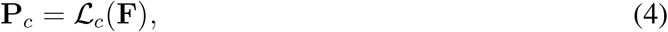

Where 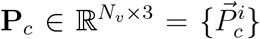, *i* = 1, 2, · · ·, *N*_*v*_, is the concatenation of the target cranial position across all *N*_*v*_ views, similarly for 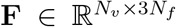. To create a general facial-to-head landmark prediction model that is applicable to a large population of subjects, we further concatenate **P**_*c*_ and **F** from all available head models, *s* = 1, 2, · · ·, *N*_*s*_, in the LYHM library, and create

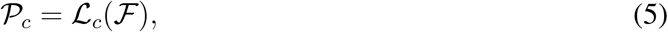

where 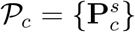 and ℱ = {**F**^*s*^}· with *s* = 1, · · ·, *N*_*s*_. Both 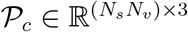 and 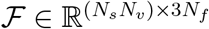 are defined in the MediaPipe coordinate system.

In determining ℒ_*c*_, we test three models: feed-forward neural network (FNN), random forest (RF), and linear transform matrix, detailed below.

#### Feed-forward neural network (FNN)

An FNN^32^ is developed, consisting of three fully connected layers, each followed by a ReLU activation function^33^ to introduce nonlinearity, and dropout layers to prevent overfitting.^34^ The model is trained with parameters including a learning rate of 0.001, a hidden dimension of 100, and dropout rates of 0.1 and 0.4.^35^ To enhance the performance of FNN, the input layer of the FNN reads not only all coordinates in ℱ, but also the pairwise distances and angles between facial landmarks.

#### Random forest (RF)

An RF model is also tested, initiated with 100 trees, with a setting of unlimited tree growth to deeply learn from the data and reduce bias.^36^ We set the split criteria^36^ so that at least two samples were required to be split, ensuring meaningful divisions. The minimum number of samples required to create a leaf node is set to one, allowing detailed pattern recognition. Bootstrapping^37^ is activated, allowing for better model generalization by training each tree on a random subset of data.

#### Linear transformation

A linear transformation offers a straightforward method to map facial landmarks to cranial landmarks. It can be expressed as

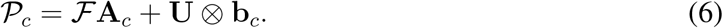

where the transformation matrix 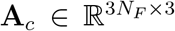, and the translation vector **b**_*c*_ ∈ ℝ^1×3^ converts facial landmarks ℱ to a specific cranial reference position 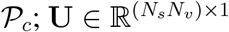 is an all-one vector, and ⊗ denotes the Kronecker product.

Based on Eq. 6, we can solve for **A**_*c*_ and **b**_*c*_ by applying a pseudo-inversion as

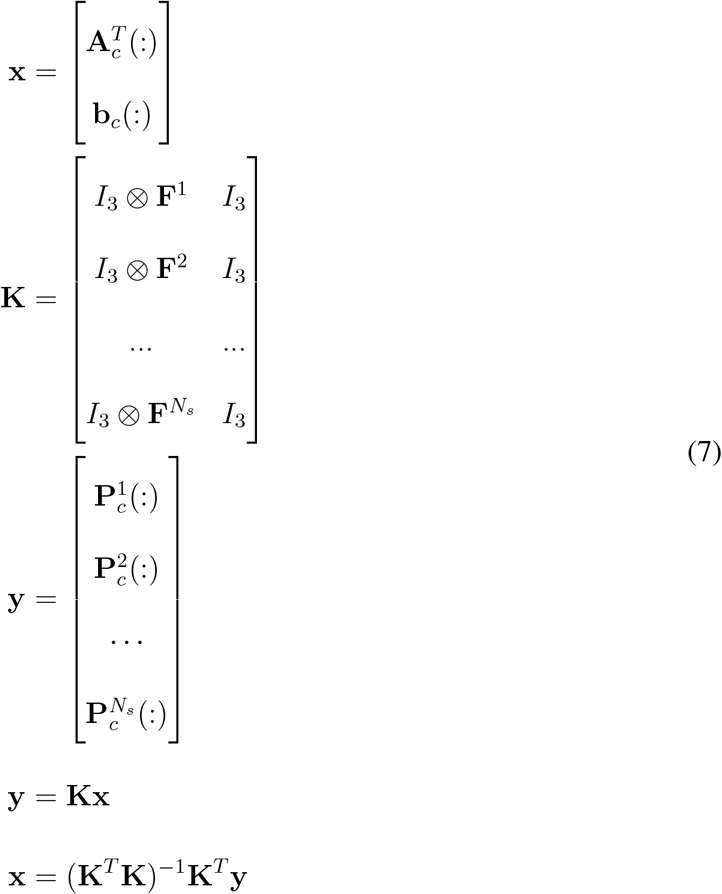

where *I*_3_ represents a 3 × 3 identity matrix. The notation “(:)” converts a matrix into a vertical vector using the column-major storage order.^38^

### 2.7 Pre-computed head landmarks from atlases and real-time rendering

To render the head 10-20 landmark positions (**H**) overlaying on top of the live camera video stream, a set of pre-computed 10-20 positions over selected atlas head model are created first (shown in Fig. 1 as shaded green). This is achieved by first manually identifying the cranial landmarks (**C**_*a*_) on a triangular surface extracted from the atlas head model and subsequently applying the Brain2Mesh toolbox^12^, ^39^ to compute the atlas 10-20 positions (**H**_*a*_).

To evaluate the impact of the chosen atlas on the accuracy of the output head landmarks, here, we compare two head models to extract 10-20 positions: an age-independent adult model based on the Colin27 atlas,^40^ and age-matched atlas models computed based on the Neurodevelopmental MRI database.^41–44^ In this study, a total of 12 age-matched atlases are used.

To register 10-20 positions **H**_*a*_ derived from atlases to an individual subject’s head shape, we use the previously obtained linear transformation model ℒ, as described in Section 2.6. We use predicted landmarks 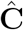 and atlas landmarks **C**_*a*_ to fit an affine transformation. This transformation is then used to map **H**_*a*_ onto the individual’s head to produce subject-specific head landmarks 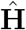.

### 2.8 Augmented reality software interface for real-time guidance

To streamline the above steps, we have developed a Python-based graphical user interface (GUI) to provide real-time guidance for placement of neuroimaging probes. A screenshot of this software is shown in Fig. 3. This GUI features two sections: a user control panel and a live video stream display. The control panel allows users to start or stop the video stream and select the preferred atlas model (including both age-independent and age-matched atlas). In addition, users can specify the density of the 10-20 systems, including 10-5, 10-10, and 10-20. In addition, the GUI also permits manual offset to the transformed 10-20 positions to further compensate for misalignments with the subject’s head contour. In this GUI, a user-configurable option is also provided to allow switching between the “3-point method”, in which LPA, RPA, Nz, Iz are coplanar, and the default “5-point method” where LPA/RPA/Nz/Iz/Cz are independently selected based on the chosen atlas. Adjustments made in the control panel are immediately applied to the registration calculations with effects shown in real time, allowing users to further fine-tune the alignment during the experiment.

**Fig 3.**
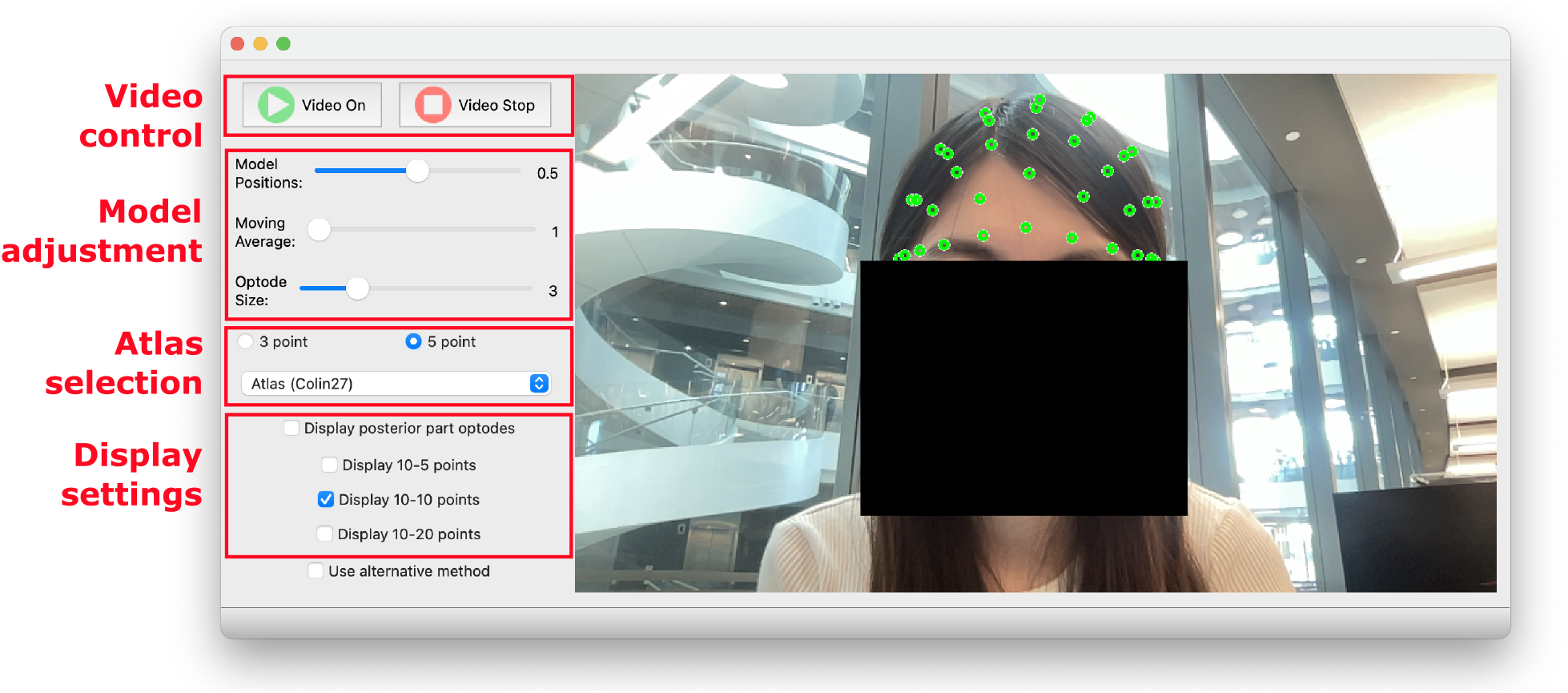
Graphical user interface (GUI) for real-time head landmark rendering and tracking in NNAR. The left panel (red boxes) shows the user control panel; the right panel (blue box) displays the video stream captured by the camera with real-time rendering of the computed 10-20 head landmarks as overlays. The face in the image is of one of the authors and is used with their consent.

The image processing workflow for each frame captured from the video stream is illustrated in the bottom row of Fig. 1. For every camera frame that contains a recognizable face, MediaPipe is used to automatically detect and compute 3-D facial landmarks. These landmarks are then transformed using the LYHM library-derived transformation matrices to compute the corresponding cranial landmarks. Subsequently, precomputed 10-20 positions based on the selected atlas model are registered to the subject’s head shape and are subsequently registered to the 2-D camera image coordinate system and displayed as overlays over the live video stream. An animation showing the real-time rendering of subject-adaptive 10-20 head landmarks is shown in Video 1.

### 2.9 Validation methods

#### 2.9.1 Validating cranial-landmark prediction

To evaluate the performance of our cranial landmark prediction models *L*, we employ a 5-fold cross-validation for each estimation technique: FNN, RF, and the linear transformation matrix, as described in Section 2.6. To achieve this goal, we randomly select 80% of the head-models available in the LYHM library for estimating *L* and subsequently test the estimated transformation using the remaining 20% of the head models. This process is repeated five times so that each head model is used once for testing and the accuracy of the prediction model is compared across different estimation techniques. This cross-validation approach is designed to minimize selection bias and enhance the reliability of our models.

#### 2.9.2 Validating 10-20 head-landmark prediction

To validate the accuracy of our head-landmark prediction models, we use the Brain2Mesh toolbox^39^ and the manually selected LPA/RPA positions, as described in Section 2.3, to calculate the reference 10-20 landmarks **H**′ as the “ground-truth”. On the other hand, the subject-specific 10-20 points 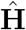 are computed by first estimating the subject cranial landmarks 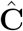 using facial landmarks, and then computing a registration to map pre-computed atlas 10-20 landmarks to the subject’s head surface. The errors between the facial-landmark derived 10-20 positions, 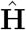 and the ground-truth 10-20 points **H** are computed to assess the accuracy of the proposed methods.

We report the overall prediction error across all tested head models, as well as the errors associated with the anterior – spanning from the coronal medial line (e.g., C3–Cz–C4) to the nasion (front of the head), and posterior – spanning from the coronal medial line to the inion (back of the head) sections of the head. The 10-20 positions along the coronal medial line are included in both regions.

### 2.9.3 Quantifying estimation noise of MediaPipe estimated facial landmarks

In our experiments, we observe that MediaPipe produces slight variations in anatomical landmarks caused by the noise in the input image. To quantify such noise-dependent landmark estimation errors, we select a sample static face image from one subject and add random Gaussian noise (mean 0 with standard deviation of 10, in units of the image pixel intensity) to create 100 testing input image sets. The choice of the noise parameter is to introduce perturbations to the image while keeping the face region detectable. By applying our aforementioned head-landmark estimation pipeline, we can quantify this image noise-induced error in both the estimated facial landmarks and the final head 10-20 position outputs.

To mitigate such noise-induced MediaPipe errors, we apply a moving average to use the mean positions averaging the landmarks estimated in successive frames to stabilize the output. We test different averaging window sizes, including 3, 5, 8, and 10 frames, to determine the trade-off between accuracy and speed.

#### 2.9.4 Assessing accuracy of real-time 10-20 prediction on human subjects

We further test our AR-based head landmark prediction pipeline on a human volunteer in a case study. In this experiment, we manually measure the subject’s head landmarks using a tape-measure, and place fiducial markers (1.9 cm diameter disk-shaped stickers) on the subject’s head at selected 10-20 locations. For each fiducial position tested, we record a video while allowing the subject to tilt head in various directions. By post-processing the recorded video, we extract the centroid of the disk-shaped fiducial marker and track it continuously using the open computer-vision (OpenCV) library; the loci of these centroid positions form a continuous trajectory (*L*_0_) in the image pixel space, serving as the ground-truth.

On the other hand, for each recorded video frame, we apply our AR workflow to predict the locations of the 10-20 positions based on the MediaPipe computed facial landmarks. The 3-D positions of the selected 10-20 locations are then projected into the pixel space of the image, forming another loci, *L*_*AR*_. Comparing the distances between the corresponding positions in *L*_0_ and *L*_*AR*_, we can quantify the accuracy and reliability of the 10-20 position estimation of our algorithm at various head tilt angles.

#### 2.9.5 Repeatability across multiple measurement sessions

In addition to assessing landmark tracking accuracy within a single session, we also repeat the above experiment across multiple measurement sessions using the same human volunteer. Again, fiducial markers were placed at two manually measured 10-20 locations – *L*_0_ and *L*_*AR*_. Following the first measurement session as described in Section 2.9.4, we repeat the same measurement in a different room and ambient lighting conditions 2 hours apart. In the first session, the room was lit with a mixed light source with overhead indoor lighting and sun light from a nearby window. In the second session, the subject is only lit by indoor lighting (overhead) in a room without a window. Variations in ambient lighting conditions, camera views, and independent data acquisition allow us to simulate realistic experimental condition differences across multiple measurement sessions. The variation in estimated 10-20 positions between the two repeated sessions was then quantified to assess the consistency and repeatability of NNAR.

#### 2.9.6 Exploring the lower-bound of 10-20 prediction error using subject-specific head surface

The linear transformation model mapping facial (**F**) to head landmarks (**H**), as detailed in Section 2.6, was derived from a large digital head model library (for estimating the face-to-head mapping) alongside with the Colin27 adult atlas (for deriving the 10-20 positions). While our underlying datasets are designed to accommodate diverse ages, genders, and head shapes, we anticipate that it may also result in suboptimal predictions for a specific subject. Here, we want to explore the lower-bound of the 10-20 prediction by minimizing the influence of subject variations and atlas biases.

To achieve this goal, we derive subject-specific linear transformations using selected subjects from the digital head model library. First, by rendering the 3-D digital head models at an additional 14 camera viewing angles, in addition to the 9 views already acquired in Section 2.1, we produce an expanded dataset including 177 subjects, each with 23 views. For each selected subject, we use a subset of 18 views (approximately 80%) to estimate the subject-specific facial-to-head landmark linear mapping and the remaining 5 views (approximately 20%) for testing. Given the limited number of acquired views, we choose to use a subset of 5 facial landmarks instead of 20 as used in ection 2.6 to predict head landmarks. In addition to capturing subject-specific images, we also pair each individual with an age-matched atlas model from the Neurodevelopmental MRI database.^41–44^ The combination of using subject’s own images to derive the linear transform for computing cranial reference points and the use of age-matching atlas for transforming cranial reference points to 10-20 landmarks are expected to provide an estimate on the lower-bound of our 10-20 prediction method.

## 3 Result

We first compare the performance of our cranial landmark prediction methods using different faceto-head transformation methods *L*. Figure 4a reports a comparison between the linear transformation, RF, and FNN methods using a dataset of 872 head models, each with 9 different viewing angles, from the LYHM library. Using the linear transformation, we found that the median errors of the predicted cranial points **P**_*c*_, where *c* ∈ {LPA, RPA, Iz, Cz}, are 0.87 cm, 0.91 cm, 1.75 cm, and 1.35 cm, respectively, resulting in an overall average error of 1.22 cm for predicting cranial landmarks. The corresponding interquartile ranges (IQRs) are 0.63 cm, 0.60 cm, 1.20 cm, and 0.94 cm, indicating relatively small errors in LPA and RPA but increased variability at Iz and Cz. Permissions were obtained for all images and videos captured from testing subjects included in this study.

**Fig 4.**
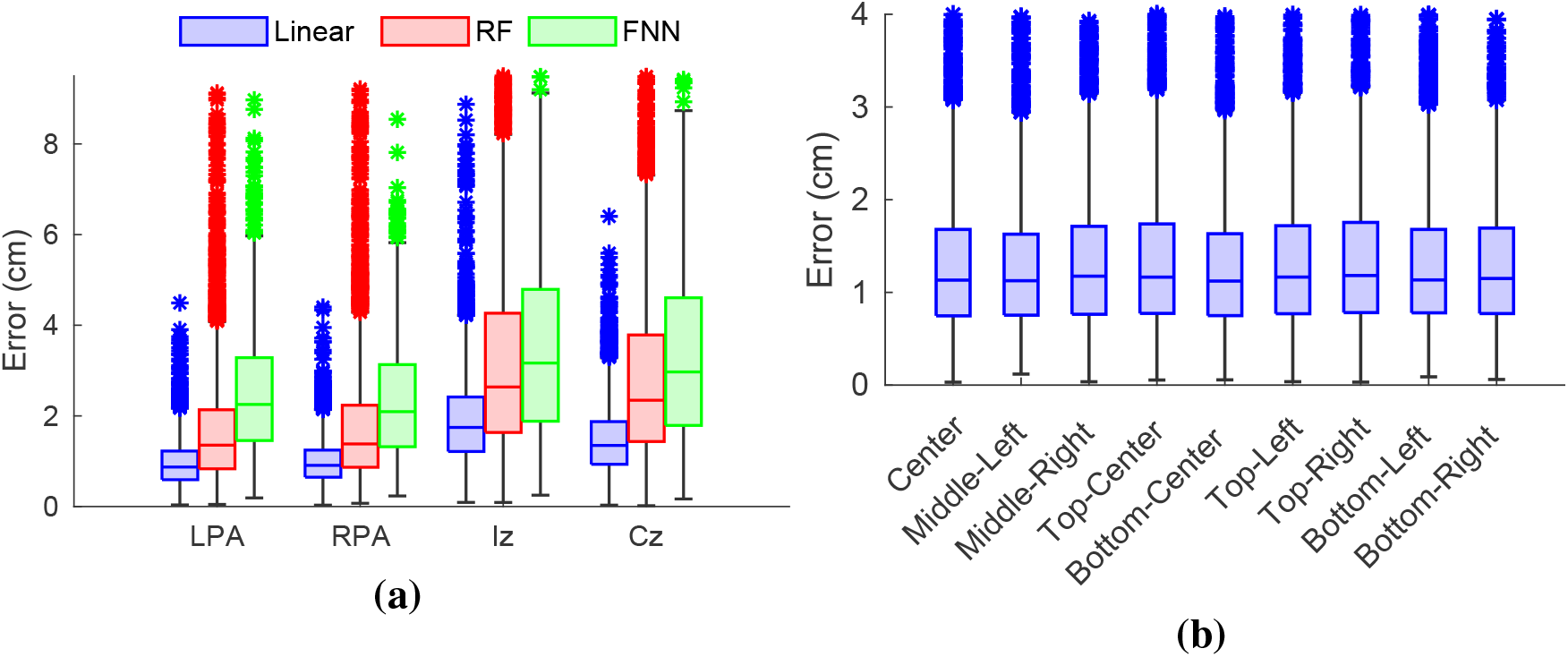
Cranial landmark prediction error assessment. In (a) we compare errors from linear transformation, random forest (RF), and feed-forward neural network (FNN) using 872 3-D digital head models, reporting error distributions at four cranial landmarks: left preauricular point (LPA), right preauricular point (RPA), inion (Iz), and vertex (Cz). In (b), we show the prediction errors of LPA/RPA/Iz/Cz across 9 different viewing angles.

When using the RF approach, the median errors are 1.35 cm for LPA, 1.38 cm for RPA, 2.64 cm for Iz, and 2.35 cm for Cz, resulting in an average prediction error of 1.93 cm. In comparison, the FNN produces median errors of 2.25 cm for LPA, 2.09 cm for RPA, 3.17 cm for Iz, and 2.97 cm for Cz, which averages to 2.62 cm across all cranial landmarks. The linear transformation approach demonstrates the lowest prediction error in all four predicted cranial landmarks. Consequently, in the remaining portion of this work, we exclusively use the linear transformation method for cranial reference point predictions.

In Fig. 4b, we report cranial landmark prediction errors using a linear transformation based on facial images captured from different viewing angles. A relatively uniform error distribution is observed, suggesting that our cranial landmark estimation approach is not sensitive to camera positions relative to the face.

Next, we evaluate the error in head landmark positions **H**. For a comprehensive analysis, we compute the estimation errors using the 10-5 system, which includes 10-10 and 10-20 as its subset. Figure 5 presents the results using 3 atlases: 1) the Colin27 atlas model,^40^ 2) the age-matched atlas model, and 3) the subject-specific head surface. Five of the 9 captured views, shown as the top-row in Fig. 2, are used in this analysis. The results revealed a median error for all 10-5 points of 1.52 cm with the Colin27 model, 1.33 cm with the age-matched model, and 0.75 cm with the head landmarks computed from subject-specific head surfaces (which eliminate inter-subject variations). Similarly to Fig. 4b, comparisons between viewing angles show little differences between the selected views (results not shown).

**Fig 5.**
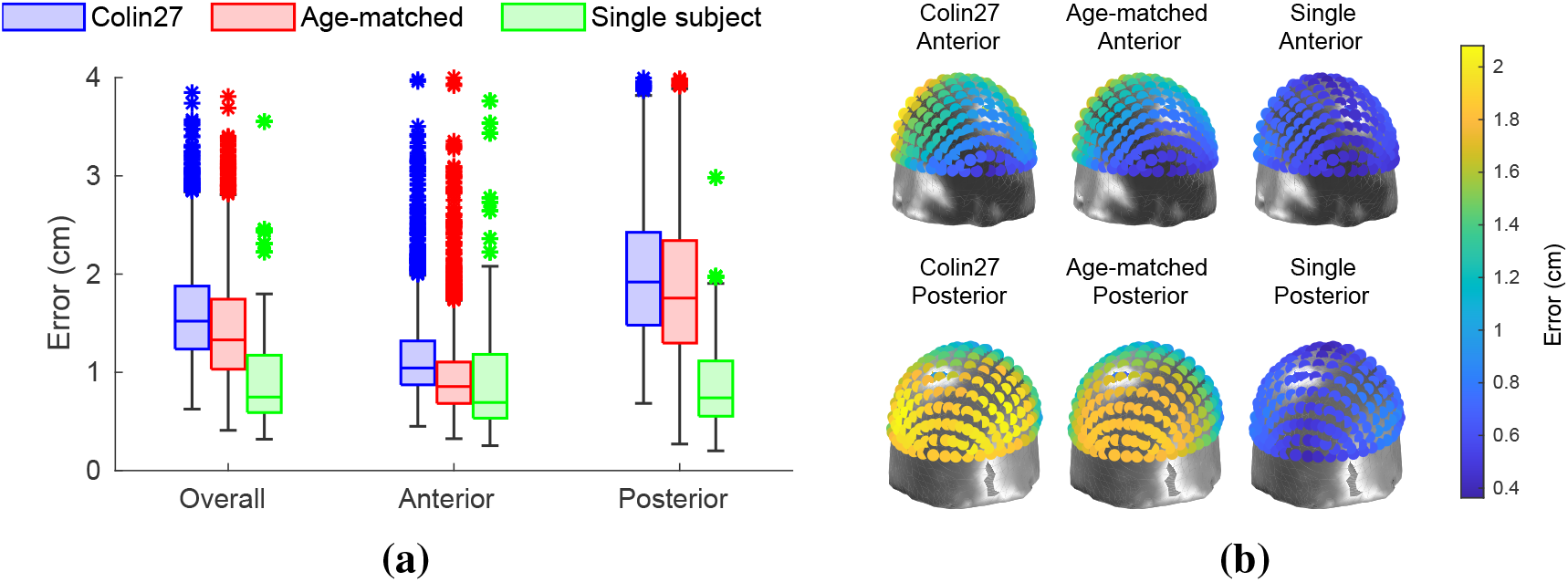
Comparison of errors in head landmark prediction over a 10-5 system using 3 different head models. Box plots in (a) illustrate the prediction errors for the 10-20 system points on the overall head, anterior head, and posterior head. Three different head models are compared: the Colin27 atlas (blue), age-matched atlases (red), and subject-specific head surface (green). The spatial distribution of median prediction errors shown in (b) are plotted along each 10-5 landmark (top: anterior view, bottom: posterior view).

We further compare the head landmark estimation errors between age and racial groups. With the Colin27 atlas, the 10-20 landmark median errors for each race group is 1.50 cm for Caucasian, 2.44 cm for Asian, 1.53 cm for African, and 1.60 cm for other origins. With age-matched atlases, the corresponding median error is 1.31 cm for Caucasian, 2.28 cm for Asian, 1.15 cm for African, and 1.35 cm for other origins. When comparing between different age groups, median errors are 1.49 cm (20s), 1.55 cm (30s), 1.56 cm (40s), 1.49 cm (50s), 1.53 cm (60s), and 1.66 cm (70s). When using the age-matched atlas model, the corresponding median errors are slightly reduced to 1.27 cm (20s), 1.37 cm (30s), 1.40 cm (40s), 1.35 cm (50s), 1.35 cm (60s), and 1.50 cm (70s). Using Welch’s analysis of variance (ANOVA) tests,^45^ we found no statistically significant differences between adult age groups in the estimation error when using Colin27 (*p* =0.40) or age-matched atlases (*p* =0.20).

Figure 5b illustrates the spatial distribution of the median prediction errors using the Colin27 atlas at all 10-5 positions tested. The errors in the posterior part of the head are larger than those in the anterior part, peaking at 2.08 cm error at the back and with a minimum of 0.41 cm error at the front. For the age-matched atlas, the maximum median error is 1.90 cm in the posterior and 0.41 cm in the anterior of the head. These are further reduced with subject-specific head surface to 0.88 cm and 0.36 cm, respectively.

The tracking and comparison between the estimated AF4 and FpZ landmark positions of a healthy volunteer are shown in Fig. 6. In this plot, the OpenCV computed trajectories (*L*_0_) of the two landmarks are shown in blue, and the corresponding AR-predicted landmark trajectories (*L*_*AR*_) are shown in orange. The distance between the landmark locations estimated from the two methods is computed in each corresponding frame, reporting an average error of 0.34 *±* 0.25 cm for AF4 and 0.11 *±* 0.08 cm for FpZ. The repeated measurement of FpZ, performed over two hours after the initial test in a different room and lighting condition, reported an average error of 0.13 *±* 0.09 cm.

**Fig 6.**
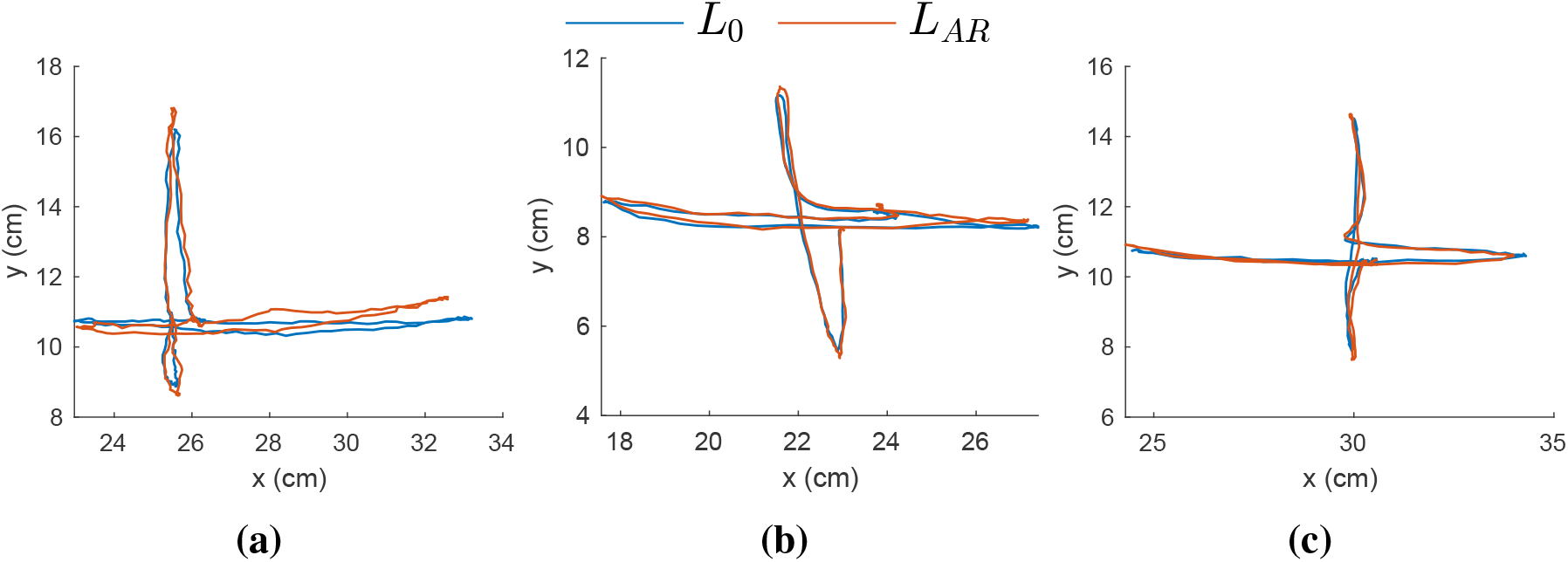
Tracing and comparison of AR-predicted 10-20 points versus manually measured 10-20 points on the forehead of a single subject. Fiducial markers are placed on two selected positions: (a) AF4, (b-c) FpZ. The FpZ tracking experiment was performed twice, shown in (b) and (c), in different room and lighting conditions for assessing the repeatability of our approach. The plots compare trajectories for AR-predicted (*L*_*AR*_, orange) and OpenCV estimated (*L*_0_, blue) AF4/FpZ locations as the head moves in four directions (top, down, left, right).

Figure 7 summarizes the noise-induced uncertainty in the 10-20 position estimated using the facial landmark data computed by MediaPipe. The maximum uncertainty (~1.71 mm), measured by the standard deviation (std), appears at the posterior part of the head (see Fig. 7a) before using averaging; this error is further reduced to 0.96 mm, 0.77 mm, 0.54 mm and 0.48 mm when using increasing moving average window length at 3, 5, 8 and 10, respectively.

**Fig 7.**
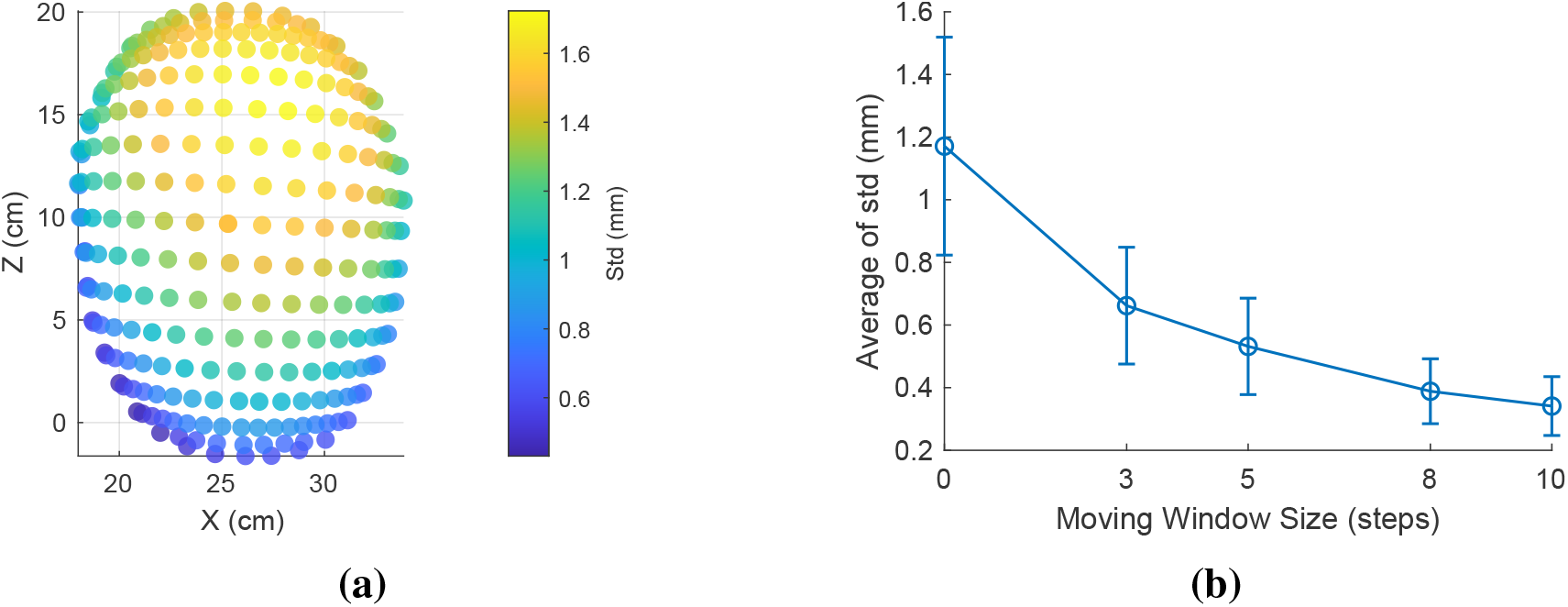
Characterization of 10-20 position errors due to the noise-induced uncertainty of MediaPipe facial position estimations of one subject. We plot in (a) the spatial maps of standard deviations of head landmark predictions from 100 noisy images was calculated to quantify the instability caused by MediaPipe. The plot in (b) illustrates the effect of applying different moving average window size (3, 5, 8, 10 steps) on the standard deviation of predictions, presenting that averaging more images reduces the standard deviation and mitigate the prediction instability issue.

## 4 Discussion

In this research, we have developed NNAR, an AR-based real-time head landmark estimation and tracking system to guide headgear placement. Using a combination of computer vision, machine learning techniques and the use of a large realistic 3-D head library, our approach offers a practical pathway to enhance headgear placement repeatability. It specifically addresses the challenges of consistent and anatomically guided probe placement, offering benefits in enhancing the data consistency between measurements in longitudinal studies and group-based analyses.

In Fig. 4a, we observe that the average of the median prediction error of the cranial landmark 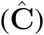 is 1.22 cm when using the linear transformation method. In comparison, other estimation methods we tested, including RF and FNN, report higher estimation errors at 1.93 cm and 2.62 cm, respectively, despite their increased complexities. We hypothesize that the added modeling complexities in RF and FNN may require more sample data in order to produce robust estimations.

In Fig. 5, we further assess the accuracy of the prediction of 10-20 head landmarks. Again, we use the manually determined cranial landmarks on the LYHM head models and subsequently compute 10-20 points over the individual 3-D head model as the ground truth. The composition of the 10-20 position estimation errors is more complex, including contributions from 1) errors resulting from facial-landmark-to-cranial landmark transformations, 2) anatomical differences between the LYHM head models used to estimate the linear transformation and the subject’s specific head shape, and 3) mismatch between the atlas head surface, over which the 10-20 positions before registration were computed, and that of the subject’s head surface. The error sources #1 and #2 are partly assessed in the results shown in Fig. 4. In Fig. 5, the 10-20 position errors are further compounded with the error source #3.

From Fig. 5, the median estimation error averaged across all 10-20 positions progressively reduces from 1.52 cm to 1.33 cm when replacing the Colin27 atlas, an age-independent adult template, by age-matched atlases, and decreases further to 0.75 cm when using the subject-specific head surface (when available). This suggests that reducing the error composition #3 could have a major impact on the accuracy of the 10-20 position estimation. The last case is considered to provide a lower-bound for the 10-20 position estimation error, because it eliminates both the error sources #2 and #3.

The results in Fig. 5 also suggest that the 10-20 position estimation error has a spatial distribution: positions close to the forehead (anterior) report the lowest estimation error, around 5 mm; the error increases when moving towards the back of the head (posterior), with a maximum error around 2 cm. This result is expected because the proposed method starts with the estimation of the facial landmarks, which are exclusively distributed over the anterior portion of the head surface. Due to the lack of imaging features towards the top of the head, as well as the obscurity of the hair, the estimation of 10-20 positions over the top of the head is largely scaled based on the cranial reference positions (LPA/RPA/Nz) estimated based on the facial landmarks, showing increased errors along the anterior-to-posterior direction. Because the back of the head is mostly invisible from the camera images, the estimation of Iz and the 10-20 positions further deviate from the ground-truth positions, resulting in the highest error over the head surface. The relatively high estimation error on the posterior part of the head is an intrinsic limitation when using a single face-directing camera.

There are a number of potential solutions to mitigate the elevated 10-20 prediction error in the posterior region of the head. A relatively simple approach is to let the operator measure the distances between Iz and Nz and provide this distance in the NNAR GUI. By scaling the NNAR estimated cranial positions using this measured distance, we anticipate that the accuracy in the posterior regions of the head can be significantly improved. Another approach is to combine head landmarks on the anterior/frontal regions of the head predicted by NNAR with a semi-rigid, 10-20 embedding headcap, such as the 3-D printable headcap published recently by our group.^12^ By aligning the headcap based on anterior landmarks (with relatively high accuracy), the embedded 10-20 positions on the headcap can provide relatively accurate head landmark guidance in the posterior regions. Furthermore, our group recently reported another machine-learning based workflow to rapidly reconstruct the subject’s 3-D head surface using real-time facial landmarks.^46^ In this work, the 3-D head surface model can achieve improved accuracy by averaging head surfaces recovered from multiple camera viewing angles. Although this approach is more computationally expensive during the head model estimation step, it offers more detailed 3-D head shapes and real-time rendering capabilities.^46^

Compared to the 0.8-2 mm estimation error reported in photogrammetry based head surface acquisition methods^47^, ^48^ and 1.7-3 mm errors when using digitizers,^17^ the estimation errors, ranging between 0.5 cm to 1.1 cm yield from our approach appear to be high. However, we want to highlight that neither photogrammetry nor a 3-D digitizer has the capability to estimate head/10-20 positions in real-time. Photogrammetry requires taking several dozen photos from various angles,^48^ and digitization requires dedicated hardware as well as careful acquisition. These methods only serve for the purpose of acquiring 3-D head shape and optode locations, and are unable to provide real-time guidance for headgear placement.

From the results of the prediction of the cranial landmarks shown in Fig. 4a, the estimation errors of Cz and Iz are higher compared to those obtained from LPA and RPA. We suspect that the reduced accuracy of Cz and Iz is the result of multiple factors. First, the MediaPipe-detected facial landmarks are closer to LPA/RPA than Cz and Iz, resulting in amplified errors at the latter locations. Second, the head surface acquired in the LYHM library contains systematic errors due to the presence of hair. In these head models, the subject’s hair is compressed and covered by a colored fabric (as shown in Fig. 2). Variation in the elevated head surfaces due to hair could impede the accuracy of Iz and Cz. Despite these challenges, our AR-based 10-20 prediction models offer real-time monitoring capability with a *<* 2 cm error (when using a linear transformation) for most 10-20 positions; those located above the frontal lobe present even lower (*<* 1 cm) error. When applying NNAR to guide probe placement, it is perceivable that aligning the head cap guided by the landmark located on the anterior portion of the head surface could yield reliable placement for the whole-head cap. The open-source GUI shown in Fig. 3 demonstrates the feasibility of real-time 10-20 position rendering, tracking, and probe placement guidance. Recognizing the limitations of our method, the GUI also permits users to manually adjust *x/y/z* offsets to allow the computed 10-20 positions to visually match the subject’s head surface.

Because of the intrinsic difficulties of estimating the Iz position using only the head surface models (such as those in the LYHM library), our cranial landmark prediction model ℱ_*c*_ can only estimate the Iz position based on the “3-point” method, where Iz, LPA, RPA and Nz are assumed to be co-planar. Despite that we were only able to validate the “3-point” method due to the lack of Iz in the used dataset, the more accurate estimation of Iz without imposing the co-planar assumption, i.e. the “5-point” method, can still be incorporated with our 10-20 prediction model. This more general anatomical assumption was not included as part of ℱ_*c*_, but as part of the predetermined cranial positions (**C**_*a*_) of the selected atlas. In our GUI, we provide options for users to toggle between these two hypotheses.

The real-time tracking capability of our AR workflow, as demonstrated in Fig. 6, also reports excellent accuracy, with an error less than 3 mm at various head tilt angles compared to the groundtruth determined by tracking the fiducial markers. The repeatability test for the FpZ landmark, shown in Fig. 6c, performed in separate rooms under different lighting conditions and camera-to-subject distances, suggests that the estimated 10-20 positions are relatively consistent for the same subject across multiple experiments. In our future studies, we hope to expand this characterization toward landmarks that are beyond the forehead area. This is currently not possible due to the presence of hair. The computation for facial landmark extraction and cranial/10-20 position prediction is quite fast, with an estimated speed of around 15 frames per second on a MacBook Air laptop with an Apple M1 processor.

The results in Fig. 7 show some intrinsic uncertainties in the results produced by MediaPipe, showing a maximum noise-induced variation of 1.71 mm when using a single image frame. However, by applying a moving average between the results obtained from subsequent image frames, we can substantially reduce this uncertainty to 0.77 mm when using an average window length of 5. The drawback of applying increasing averaging is a noticeable lag in the predicted positions due to the use of older positions. In the GUI, we allow users to set the averaging window length to balance between such lag and the accuracy. Because the overall uncertainty from MediaPipe is relatively low, around 1 mm even without averaging, all the results presented in this manuscript were obtained without applying averaging. However, as shown in Fig. 7, when additional accuracy is desired, averaging can further reduce the error. We also note that in certain experiments, averaging could introduce lag, which could be an issue for subjects with head movement.

We would like to mention a few limitations of this study. First, using a single linear mapping fitted from a large group of subjects with varying demographics and head sizes inevitably introduces a trade-off between accuracy and generality. This limitation is particularly responsible for the nearly doubled 10-20 estimation error (2.44 cm) in Asian head models compared to those from Caucasian origins (1.50 cm). Because 92.7% of the head models in the LYHM library were obtained from Caucasian subjects and Asian head models only account for 3% of the used dataset, the increased prediction errors suggest that increasing the sample size and deriving separate face-to-head mapping transformations for each racial group are important for further improvement of this work. Interestingly, no statistically significant differences were found between various adult-only age-groups. Extending these characterizations to newborns and youth populations could expand the use of NNAR in studies involving pediatric subjects. Using machine learning-based models was expected to offer better generality towards racial and age differences due to the increased number of parameters. However, the simple machine learning-based approaches we have tested, such as RF and FNN, were not effective. Investigating more advanced machine learning-based faciallandmark-to-head mapping networks could provide pathways to further improve the accuracy of this work in the future.

Moreover, the use of facial landmarks requires the camera image to present a good coverage of the subject’s face. Views of the camera from the side of the subject or partial face coverage would render this approach ineffective. Additionally, the error plots in Fig. 5 exhibit a consistent pattern, indicating that this approach produces better accuracy for the anterior part of the head compared to the posterior part of the head surface. We believe this is because our 10-20 positions are computed using linear mappings based on facial landmarks, thus errors scale up when moving away from facial landmarks. Methods utilizing multiple viewing angles can be developed to circumvent this limitation. Finally, our workflow was built upon MediaPipe – an widely used open-source facial detection framework. Based on the results in Fig. 7, we have identified MediaPipe-induced uncertainties, despite being small, which generally align with the characterization of errors reported by the MediaPipe team,^25^ among others.^49–51^ Exploring more robust facial landmark processing tools could be an important future step for improving accuracy.

## 5 Conclusion

In summary, in this work, we have developed NNAR, an AR-based workflow to estimate, render, and continuously track 3-D cranial landmarks and head 10-20 landmarks in real-time, offering opportunities for guiding consistent fNIRS/EEG headgear placement during experiments, as well as potentials for shortening the headgear setup time. The cranial and head 10-20 landmarks are computed using facial landmarks based on registration methods derived from a large high-quality head surface library and brain atlases. This subject-agonistic registration can readily accommodate diverse subjects of different ages and gender groups without needing additional inputs or measurements, however, the accuracy when using the subject-agonistic registration is moderate, reporting a *<* 1 cm median error for landmarks over the anterior part of the head, and *<* 2 cm errors for landmarks over the posterior part of the head surface. We also found that by replacing the subject-agonistic registration with that derived from age-dependent atlases or subject-specific head surfaces, if available, one can reduce the median position errors of 10-20 positions roughly by half. Finally, we have implemented this AR approach in NNAR, an easy-to-use open-source GUI, enabling rapid dissemination across broad fNIRS/EEG research communities.

## Disclosures

No conflicts of interest, financial or otherwise, are declared by the authors.

## Code, Data, and Materials Availability

The source code for the AR 10-20 monitoring software can be downloaded at https://github.com/COTILab/NeuroNavigatAR; a library of facial and cranial landmarks derived from the LYHM library is available at https://neurojson.org/db/cotilab/NeuroNavigatAR.

## Acknowledgments

This research is supported by the National Institute of Health (NIH) National Institute of Biomedical Imaging and Bioengineering grant R01-EB026998 and the National Institute of Neurological Disorders and Stroke (NINDS) grant U24-NS124027. We thank Dr. Yi-han Sheu for insightful comments that contributed to refining our methodology.

## Notes

### Competing Interest Statement

The authors have declared no competing interest.

